# Improvement of *Neisseria gonorrhoeae* culture media to enable growth without CO_2_

**DOI:** 10.1101/2023.08.01.551449

**Authors:** Chukwuma Jude Menkiti, Lori A. S. Snyder

## Abstract

Bacterial culture is the crucial step in diagnosing *Neisseria gonorrhoeae* infections and is the gold standard for determining their antimicrobial resistance profile. However, culture of *Neisseria* spp. can be challenging in resource poor areas, relying on specialist incubators supplying 5% CO_2_ for growth of the bacteria. Even when such incubators are available, the CO_2_ to run them may be scarce; there were CO_2_ shortages during the COVID-19 pandemic, for example. Although culture jars with gas packs or candles can be used, these are inefficient in terms of use of incubator space and researcher time. To achieve simplicity in culturing of *N. gonorrhoeae*, the standard Oxoid GC base medium, made with the Kellogg’s glucose and iron supplements was improved with the addition of 0.75 g/L sodium bicarbonate (NaHCO_3_), which is inexpensive and really available. This improved media was able to sustain gonococcal growth as well as standard GC media in 5% CO_2_. Chocolate agar and Thayer-Martin agar with sodium bicarbonate was also developed, with all showing good growth of *N. gonorrhoeae* without the need for atmospheric CO_2_.

## Introduction

*Neisseria* species are fastidious organisms and therefore require additional nutritional requirements in the form of supplementary iron (1/1000 v/v) and a glucose-based (1/100 v/v/) supplement (1) to be added to the base GC media to grow (2). The optimum temperature for the growth of *Neisseria* spp. is 35 to 37°C (3,4) and they grow best at a pH of 7 to 7.5 (2,5).

*Neisseria* spp. are carboxyphilic (capnophilic) bacteria; they require an atmosphere composed of oxygen and a raised level of CO_2_ and therefore grow best in an atmosphere enriched with 5 - 10% CO_2_ (6,7,8,9). This atmosphere can be provided using a CO_2_ incubator, which adds CO_2_ gas via a regulator, or by using a CO_2_ generating GasPak kit or candle in a culture jar. Marble chips can also be used to provide CO_2_ enriched atmosphere (2, 3, 7).

To enable laboratories in resource poor areas to be able to grow *N. gonorrhoeae* as they would other pathogens opens up prospects for wider coverage of diagnostic and antimicrobial susceptibility tests. This would mean a significant improvement in patient care and the potential for treatments to be personalized and targeted. A crucial step in achieving this is finding an alternative to the 5% CO_2_ needed for the incubation and growth of *N. gonorrhoeae*. This has been achieved by modifying the GC medium (Oxoid) with Kellogg’s supplements (1) using 0.75 g/L sodium bicarbonate (NAHCO_3_), a carbonic acid with alkalinizing and electrolyte replacement properties.

Three kinds of bacterial culture media were used in these investigations: GC; chocolate; and Thayer-Martin media.

GC agar is a media used for the cultivation of *N. gonorrhoeae* in the research laboratory. Growth supplements like peptone, iron, and glucose are included in GC media to provide nutrients needed for growth. GC media also contains cornstarch, which absorbs toxic metabolites. The base GC media can be enriched by addition of other supplements like lysed red blood cells and can be made selective through the addition of antibiotics (10,11,12,13,14,15,16). Colonies of *N. gonorrhoeae* grown on GC agar show a few different morphologies, which are the result of phase variation (17). Colonies may be transparent, appearing blue, or opaque, appearing yellow, when viewed with indirect lighting under a dissecting microscope. This is due to lack of expression and expression of Opa proteins, respectively (18,19,20). Colonies may also differ in size, smooth or rough appearance, and distinct or ruffled edges, all determined by expression or not of gonococcal pili (21). These characteristics are best observed on GC agar, which is light in colour and translucent.

Chocolate agar is an enriched media made from a GC agar base and lysed red blood cells, which gives the agar its characteristic chocolate brown colour and hence its name. The red blood cells are lysed by heating at 80°C before adding to the media (22,23,24,25). Chocolate agar includes X factor (hemin) and V factor (nicotinamide adenine dinucleotide; NAD), a coenzyme (26,27). The X and V factors from the lysed red blood cells are needed for the successful growth of fastidious bacteria including *N. gonorrhoeae* (28). Chocolate agar also contains casein, which provides the amino acids, nitrogen, and other essential nutrients and elements necessary for growth. As for GC media, the cornstarch present neutralizes metabolites that maybe toxic to highly sensitive *Neisseria* species. The presence of potassium phosphate and sodium chloride in chocolate agar maintains the pH and osmotic equilibrium of the media, respectively. These factors help maintain the integrity of the bacterial cells during growth. *N. gonorrhoeae* colony growth on GC agar appears as large, smooth, round, convex, and opaque colonies with defined edges (29).

Thayer-Martin media is a modified chocolate agar, with a GC agar base and other supplements included in the media that enhance the growth of *N. gonorrhoeae* (30,31). Unlike GC and chocolate media, Thayer-Martin media is selective, meaning that *Neisseria* spp. will grow on it, but few other bacterial species will (31,32,33,34). This is in large part due to the inclusion in the Thayer-Martin media of antimicrobials like colistin, vancomycin, trimethoprim, and nystatin (31,34). Colistin, a polymixin E, is used to inhibit the growth of Gram negative bacteria that may be plated alongside *N. gonorrhoeae*. Vancomycin is likewise used to inhibit Gram positive bacterial floras. Trimethoprim is also a very useful addition; it prevents the swarming of *Proteus* species and inhibits both Gram positive and Gram negative aerobic bacterial floras. Nystatin is an antifungal preventing the growth of yeasts (32,33,35,36).

Given the constraints of atmospheric requirements by *N. gonorrhoeae* for laboratory growth, research and development of an alternative was pursued. Ultimately, a growth media for culturing *N. gonorrhoeae* without the need for atmospheric CO_2_ supplementation will be useful for laboratory research and diagnostics. This is especially important in resource poor areas, where CO_2_ may not be readily available or in short supply.

## Materials and Methods

### GC medium

The GC medium (Oxoid) containing Kellogg’s glucose and 5% iron supplements (1) was used for the culture and isolation of all *Neisseria* spp. used in these investigations. GC media was prepared as follows: 36 g of the Oxoid GC agar base were added to 1 litre of distilled water and was sterilised at 121°C for 15 minutes in an autoclave. The media was then allowed to cool, but not solidify, before adding 10 ml of Kellogg’s glucose supplement and 1 ml of Kellogg’s iron supplement (1).

For the modified GC medium, 0.75 g/l of sodium bicarbonate was added after adding the glucose and iron supplements. In both cases, the media was thoroughly mixed and approximately 20 ml of the media was poured into 90 mm sterile petri dishes. These agar plates, both GC and modified GC, were allowed to cool and solidify. Labelled agar plates were stored at 4° and allowed to return to room temperature before use.

Kellogg’s glucose supplement was prepared using 40 g of D-Glucose, dissolved in 70 ml of distilled water (1). To solubilize the sugar, this mixture was stirred on a hot plate until the glucose was well dissolved. The glucose mixture was then cooled to room temperature before the addition of 1 g of L-Glutamine and 2 mg of Co-carboxylase (Thiamine pyrophosphate). The final volume of the supplement preparation was made up to 100 ml (1). Finally, the Kellogg’s glucose supplement was filter sterilized into sterile tubes, labelled, and stored at 4°C.

Kellogg’s Iron supplement contains 2 g Fe(NO_3_)_3_ dissolved in 40 ml distilled water. This media supplement was then filter sterilised into a sterile tube, labelled, and stored at 4°C.

### Chocolate agar

To make chocolate agar, 36 g of the Oxoid GC agar base were added to 500 ml of distilled water, according to manufacturer’s instructions. A 2% haemoglobin supplement was prepared by dissolving 10 g of Thermo Scientific™ Oxoid™ Hemoglobin Soluble Powder in 500 ml of distilled water. The haemoglobin powder was dissolved to avoid the formation of clumps. The GC agar base and haemoglobin solution were sterilised by autoclave at 121°C for 15 minutes. After sterilization, the GC agar and haemoglobin solution were allowed to cool. The sterile haemoglobin solution was then added to the GC agar base and mixed gently to avoid formation of clumps and bubbles. BD BBL™ IsoVitaleX™ Enrichment supplement was reconstituted by adding 10 ml of the accompanying diluents, mixed by shaking, then added immediately to the GC agar base / haemoglobin solution. Also added were 15 g/L Tryptic Soy Broth (BD Bacto™), 1 g/L 1-Allyl-3-methylimidazolium chloride (Alfa Aesar™), 5 g/L Sodium Chloride (Fisher BioReagents), 4 g/L Potassium Phosphate Dibasic (Fisher BioReagents), and 1 g/L Potassium Phosphate Monobasic (Fisher BioReagents). This media was gently mixed to avoid formation of bubbles, then 20 ml was poured into 90 mm sterile petri dishes. After cooling and solidifying, the plates were labelled and stored at 4°C. For the modified Chocolate medium, 0.75 g/L of sodium bicarbonate was added along with the other chocolate agar supplements.

### Thayer-Martin medium

For Thayer-Martin media, 36 g of GC agar base (Oxoid) was added to 500 ml of distilled water. Separately, 2% of haemoglobin was prepared by dissolving 10 g of Haemoglobin Soluble Powder (Oxoid) in 500 ml of distilled water and allowed to dissolve to avoid formation of clumps. The 500 ml GC agar base and 500 ml 2% haemoglobin solution were sterilised at 121°C for 15 minutes. After autoclaving and cooling, the sterile haemoglobin solution was added to the sterile GC agar base and mixed gently to avoid formation of clumps and bubbles. IsoVitaleX™ Enrichment (BD BBL™) was reconstituted by adding 10 ml of the accompanying diluents, shaking to mix, and added immediately to the media. Also added to the media was 3 μg/L Vancomycin Supplement (Oxoid), 7.5 μg/L Colistin sulfate salt (ACROS Organics™), 12.5 μg/L Nystatin (MP Biomedicals™) and 5.0 μg/L Trimethoprim lactate (Alfa Aesar). The media was mixed gently and 20 ml poured into 90 mm sterile petri dishes. Once solidified, these plates were stored at 4°C. For the modified Thayer-Martin medium, 0.75 g/l of sodium bicarbonate was added along with the supplements.

#### *Neisseria* spp. isolates

Stocks of the *Neisseria* spp. used were stored at -80°C. Freezer stocks were generated using GC broth (3.75 g proteose peptone, 1 g potassium phosphate dibasic K2HPO4, 0.25 g potassium phosphate monobasic KH_2_PO_4_, 1.25 g NaCl). The reagents were added to 250 ml distilled water, dissolved by heating and sterilised by autoclaving at 121°C for 15 minutes. Once cool to the touch, the glucose (2.5 ml) and iron (250 μl) supplements are added along with sterile glycerol to make a final concentration of 15% glycerol in GC agar. Culture samples were resuspended in at least 1 ml of this GC freeze media in Thermo fisher Fisherbrand™ Threaded Sterile Cryogenic vials for storage at -80°C.

Five isolates were used for this study: four commensal, non-pathogenic isolates and the pathogenic control, *N. gonorrhoeae* strain NCCP11945 (37). The commensal isolates used were *Neisseria subflava* isolates KU1003-01, KU1003-02, and RH3002vg and *Neisseria cinerea* isolate RH3002vg (38).

Identification tests were done throughout the experiments, to verify the *Neisseria* spp. cultures had not become contaminated, including verification of colonial morphology, Gram staining, catalase, and oxidase testing.

### Colonial morphology

On GC agar, *N. gonorrhoeae* strain NCCP11945 colonies appeared as small, round, and convex unpigmented colonies. *N. subflava* isolate KU1003-01 appeared as large, round, smooth, moist, and yellow glistening colonies. *N. subflava* isolate KU1003-02 appeared as medium, round, yellow colonies with a rough surface. *N. subflava* isolate RH3002v2g had medium, round, smooth, and yellow colonies with a glistening surface. *N. cinerea* isolate RH3002v2f were small, round, and unpigmented colonies with glistening surface.

On Chocolate agar and Thayer-Martin agar, the colonies from each of these were difficult to differentiate. After 48 hours of incubation, all of these *Neisseria* appeared as small, convex, colourless, and moist colonies with glistening surfaces.

### Gram stain

Colonies were smeared and heat fixed to microscope slides for Gram staining (39). A 1% aqueous solution of crystal violet was used to stain the smear for 1 minute. The stain is washed off with water and mordant with Lugol’s Iodine was added for 1 minute. The mordant was washed off and the smear decolourized with acetone. The acetone was washed with water and the counterstain, a 0.5% aqueous solution of safranin O was added. After 1 minute, the counterstain was washed with water, blotted dry and examined with via light microscope (Nikon Eclipse) with oil immersion. All of the *Neisseria* spp. investigated here appeared as Gram negative diplococci under microscopy.

### Catalase Test

To assess the bacteria for catalase activity (40), a clean slide was placed in a petri dish and a few colonies of the organism were emulsified in distilled water. Two drops of hydrogen peroxide (H_2_O_2_) were added to the emulsification. The appearance of bubbles indicates a positive reaction. *Neisseria* spp. are catalase positive (except *N. elongata* which is catalase negative); they produce the enzyme catalase which acts as a catalyst in the breakdown of H_2_O_2_ to water and oxygen, resulting in bubbles (4). All *Neisseria* spp. investigated here demonstrate catalase activity.

### Oxidase Test

Oxidase testing for the expression of bacterial cytochrome oxidase was used in combination with the Gram stain and catalase results to verify the identity of the growth on the media plates as being *Neisseria* spp., which are oxidase positive (4). Bioconnections CK3520 oxidase test strips were smeared with the colonies. A deep blue/purple colour developing within 30 seconds indicates a positive oxidase test (41). The oxidase strip is impregnated with a redox dye, 1% tetramethyl-paraphenylene diamine dihydrochloride (TMPPDH), which is oxidised to a deep blue/purple hue by the oxidase enzyme that is produced by some aerobic bacteria during their respiratory oxidation mechanism, including all *Neisseria* spp. in this study.

#### *Neisseria* spp. Culture Media Investigations

For the study, twelve agar plates (6 standard and 6 modified) were used for each isolate. To assess any potential and variations in bacterial growth, 3 standard and 3 modified inoculated plates where incubated a 5% CO_2_ incubator. Under this atmospheric condition, it is expected *Neisseria* spp. will grow (4, 42). The other 6 plates, 3 standard and 3 modified, were inoculated and placed in a standard incubator, being provided with a normal atmosphere. The use of 3 plates for each condition produced technical replicates. In addition, the experiment was repeated three times on different days, generating biological replicates.

Controls were used in all the experiments and were incubated alongside the media being investigated. A standard GC agar plate inoculated with *N. gonorrhoeae* strain NCCP11945 was used as control; this gonococcal strain is known to require CO_2_ for growth on GC media (37).

The same principles and protocols were also applied to Chocolate agar and Thayer-Martin agar and the sodium bicarbonate modified versions of these.

## Results and Discussion GC culture

In our quest to develop an alternative *N. gonorrhoeae* diagnostic test for laboratories, especially those in resource poor areas, bacterial culture is necessary so that antimicrobial susceptibilities can be determined and patient treatment can be personalized and targeted. Although molecular testing can indicate that a bacterial species is present and advances may be able to accurately determine if antimicrobial resistance genes are present, only a bacterial culture can verify viable infectious agents and determine expression of antimicrobial resistance.

A crucial step in achieving a wider capacity for *N. gonorrhoeae* culturing in poor areas is finding an alternative to the 5% CO_2_ needed for the incubation and growth of the bacteria. This has been achieved by modifying the GC medium (Oxoid) with Kellogg’s supplements (Kellogg *et al*., 1963) using 0.75 g/L sodium bicarbonate (NaHCO_3_), a carbonic acid with alkalinizing and electrolyte replacement properties.

There was no growth of the *Neisseria* spp. isolates on the standard, unmodified GC media incubated in the standard atmosphere incubator. All of the *Neisseria* spp. isolates were able to grow heavily on the NaHCO_3_ modified GC media in the standard atmosphere incubator and on both types of media in the CO_2_ incubator (Table 1).

**Table 1.** *Neisseria* spp. growth on NaHCO_3_ modified GC agar.

|  | Growth in CO<sub>2</sub> incubator<sup>^</sup> |  | Growth in standard incubator<sup>^</sup> |  |
| --- | --- | --- | --- | --- |
| ***N. gonorrhoeae*** | **24 hrs** | **48 hrs** | **24 hrs** | **48 hrs** |
| GC agar | +++ | +++ | - | - |
| NaHCO<sub>3</sub> modified GC agar | +++ | +++ | +++ | +++ |
| ***N. subflava*<sup>\*</sup>** | **24 hrs** | **48 hrs** | **24 hrs** | **48 hrs** |
| GC agar | +++ | +++ | - | - |
| NaHCO<sub>3</sub> modified GC agar | +++ | +++ | +++ | +++ |
| ***N. cinerea*** | **24 hrs** | **48 hrs** | **24 hrs** | **48 hrs** |
| GC agar | +++ | +++ | - | - |
| NaHCO<sub>3</sub> modified GC agar | +++ | +++ | +++ | +++ |
| * Data was the same for all three strains of *N. subflava* investigated. |  |  |  |  |
| <sup>^</sup> Growth: - No growth; + Scanty growth; ++ Moderate growth; +++ Heavy growth |  |  |  |  |

To confirm whether this alternative media was able to support growth as well as standard GC media with Kellogg’s supplements, both types of media were incubated in a CO_2_ incubator and the results compared. The *N. gonorrhoeae, N. subflava*, and *N. cinerea* grew equally well at 37°C in all three conditions: standard GC media with 5% CO_2_; modified GC media with sodium bicarbonate and 5% CO_2_; and modified GC media under standard atmospheric conditions. The experiments were repeated three times for reproducibility (Table 1).

### Chocolate agar

*Neisseria* spp. are also often grown in diagnostic laboratories on Chocolate agar, therefore it was desirable to develop a modification that would enable culture on this media without CO_2_ as well. This was achieved by modifying the chocolate agar through the addition of 0.75 g/L sodium bicarbonate, as had been achieved for GC agar.

After 24 hours of incubation, there was heavy growth of the *Neisseria* spp. on all the inoculated media incubated in the CO_2_ incubator. For the media incubated in the standard incubator, there was only growth on the NaHCO3 modified Chocolate agar (Table 2). There were moderate growth at 24 hrs of incubation and heavy growth at 48 hrs of incubation on the modified Chocolate agar plates incubated in the standard atmosphere incubator. Heavier growth could be achieved at 24hrs by sealing the modified Chocolate agar plates in a plastic bag. The experiments were repeated three times (Table 2).

**Table 2.** *Neisseria* spp. growth on NaHCO_3_ modified Chocolate agar.

|  | Growth in CO<sub>2</sub> incubator<sup>^</sup> |  | Growth in standard incubator<sup>^</sup> |  |
| --- | --- | --- | --- | --- |
| ***N. gonorrhoeae*** | **24 hrs** | **48 hrs** | **24 hrs** | **48 hrs** |
| Chocolate agar | +++ | +++ | - | - |
| NaHCO<sub>3</sub> modified Chocolate agar | +++ | +++ | ++ | +++ |
| Modified Chocolate, sealed bag | +++ | +++ | +++ | +++ |
| ***N. subflava*\*** | **24 hrs** | **48 hrs** | **24 hrs** | **48 hrs** |
| Chocolate agar | +++ | +++ | - | - |
| NaHCO<sub>3</sub> modified Chocolate agar | +++ | +++ | ++ | +++ |
| Modified Chocolate, sealed bag | +++ | +++ | +++ | +++ |
| ***N. cinerea*** | **24 hrs** | **48 hrs** | **24 hrs** | **48 hrs** |
| Chocolate agar | +++ | +++ | - | - |
| NaHCO<sub>3</sub> modified Chocolate agar | +++ | +++ | ++ | +++ |
| Modified Chocolate, sealed bag | +++ | +++ | +++ | +++ |

\* Data was the same for all three strains of *N. subflava* investigated.
<sup>^</sup> Growth: - No growth; + Scanty growth; ++ Moderate growth; +++ Heavy growth

### Thayer-Martin Agar

Thayer-Martin agar is the only media investigated here that is selective for *Neisseria* spp., including inhibitors for growth of other Gram-negative species, Gram-positive species, and fungi. As for GC agar and Chocolate agar, Thayer-Martin agar was modified through the addition of 0.75 g/L sodium bicarbonate.

As expected, there was heavy growth on all of the inoculated Thayer-Martin media incubated in the CO_2_ incubator. For the media incubated in the standard atmosphere incubator, there was only growth on the modified Thayer-Martin agar (Table 3), as seen for the other two media. Similarly to the Chocolate agar results, there was only moderate growth at 24 hrs on the modified Thayer-Martin agar in the standard incubator; this could be improved by sealing the plate in a bag (Table 3). These experiments were repeated three times.

**Table 3.**
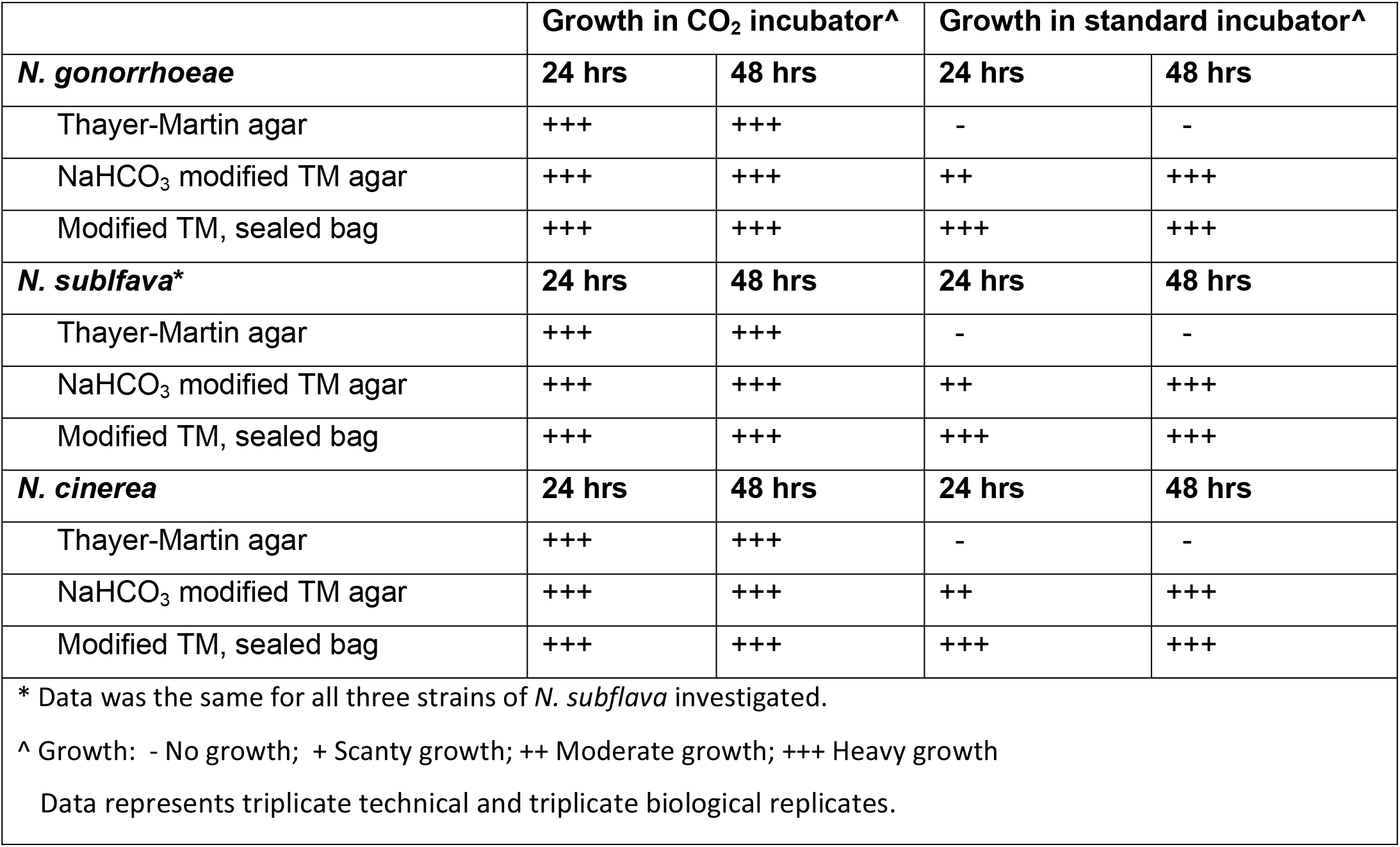
*Neisseria* spp. growth on NaHCO_3_ modified Thayer-Martin (TM) agar.

### Principle

These results demonstrate that the addition of sodium bicarbonate to GC agar, Chocolate agar, and Thayer-Martin agar provides the supplementary CO_2_ required for the growth of *Neisseria* spp. Hence, sodium bicarbonate modified media is an alternative to the use of CO_2_ incubators, candle jars, or CO_2_ GasPaks when culturing *N. gonorrhoeae* and other *Neisseria* species.

Sodium bicarbonate (NaHCO_3_), commonly known as baking soda, is a monosodium salt of carbonic acid with alkalinizing and electrolyte replacement properties. NaHCO_3_ is water soluble, with two sodium bicarbonates dissociating to form two sodium ions (2Na^+^) and two carbonate ions (2HCO_3_^-^). These carbonate ions decompose to release CO_2_ and H_2_O (Figure 1). Due to the properties of the media investigated here, the growth conditions are buffered to pH 7.2. The pKa of carbonic acid (H_2_CO_3_) is around 6.4, therefore the sodium bicarbonate will be in equilibrium within the media, readily converting into CO_2_ during culture. Because the media is buffered and a small amount of sodium bicarbonate is added (0.75 g/L), the pH of the media does not shift from the pH of 7 - 7.5 required for *N. gonorrhoeae* growth.

**Figure 1.**
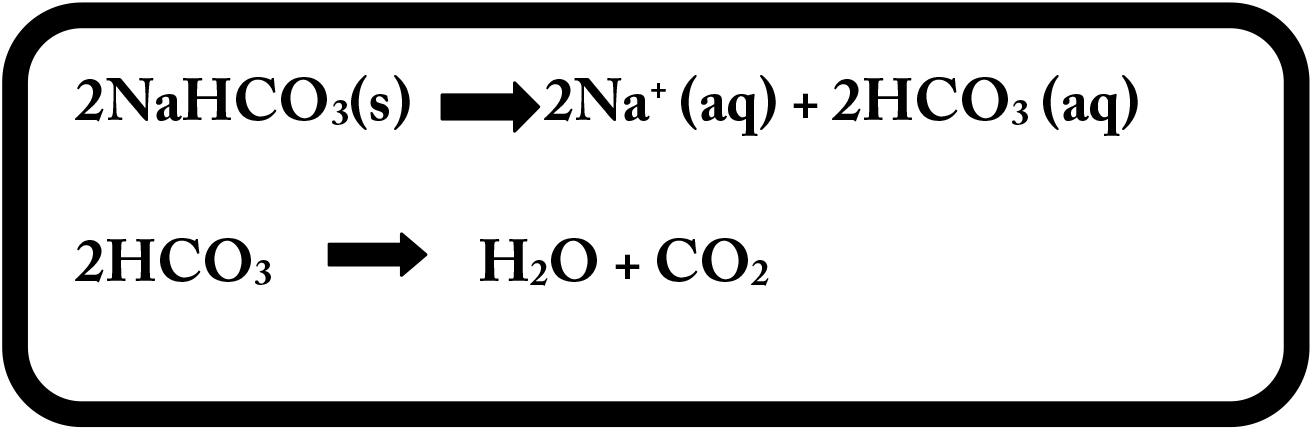
Sodium bicarbonate (NaHCO_3_), in the presence of water decomposes into sodium ions (Na^+^) and carbonate ions (HCO_3_^-^). Once the carbonate ions are formed, carbon dioxide (CO_2_) is spontaneously released. (s) Soluble compound. (aq) aqueous (dissolved in water).

## Conclusion

In the absence of enhanced levels of CO_2_, normally required for laboratory growth of *N. gonorrhoeae, N. subflava, N. cinerea*, and other *Neisseria* spp., addition of 0.75 g/L to GC media, Chocolate agar, and Thayer-Martin agar is able to support culturing. This advance in neisserial bacterial growth media alternatives opens up the prospect for more widespread research and diagnostics.

